# CLEAR-AF: Improved Autofluorescence Subtraction for Multiplexed Tissue Imaging via Polar Transformation and Gaussian Mixture Models

**DOI:** 10.1101/2025.11.22.689928

**Authors:** Alexandre Arnould, Bart De Moor, Frederik De Smet, Francesca Maria Bosisio, Asier Antoranz

## Abstract

Cyclic Multiplexed ImmunoFluorescence (cMIF) can profile dozens to hundreds of proteins at subcellular resolution, offering unprecedented insights into tissue architecture in health and disease. However, measured signals are affected by tissue autofluorescence (AF), which can confound molecular quantifications at the single-cell level. Existing computational strategies to subtract AF are not sensitive nor specific enough and often leave residual background or over subtract weak markers.

To address current limitations, we introduce CLEAR-AF (Coordinate-transformed Local Estimation and Adaptive Removal of AutoFluorescence), a subtraction framework where AF is calibrated for each acquired signal using a reference image. CLEAR-AF maps these intensities into polar coordinates to disentangle AF from true signal, and applies adaptive, distribution-aware thresholding to estimate and remove AF locally.

Across multiple cMIF technologies, CLEAR-AF yields cleaner marker channels with improved specificity, greater sensitivity, and higher reproducibility relative to common AF removal approaches. By delivering more accurate per-cell measurements without workflow changes to acquisition, CLEAR-AF provides a practical, platform-agnostic step towards more reliable spatial proteomics at scale.

## Introduction

Cyclic Multiplexed Immunofluorescence (cMIF) enables the spatial profiling of hundreds of proteins at subcellular resolution, providing unprecedented insights into tissue composition and architecture (Schapiro et al., 2021), (Lin et al., 2015), (Karimi, 2024). By simultaneously capturing molecular expression and spatial context, this technology has become central to oncology, immunology, and neuroscience, where the microenvironment strongly influences cell behaviour and therapeutic response (Antoranz et al., 2022), (Eng et al., 2025), (Zhang et al., 2025).

A fundamental challenge in cMIF arises from autofluorescence (AF), background signal emitted by endogenous tissue components such as collagen, elastin, NAD(P)H, and lipofuscin (Monici, 2005), (Lakowicz, 2006). Even in the absence of staining, specific excitation wavelengths induce fluorescent signals that spectrally overlap with fluorophores. In contrast to true signal (TS) originating from labelled protein targets, AF introduces non-specific background that biases per-pixel intensities and confounds single-cell quantifications. This problem is particularly acute in intrinsically highly fluorescent tissues (e.g., liver, lung, brain), where weakly expressed markers can become indistinguishable from background (Croce et al., 2018), (Jun et al., 2017), (Tan et al., 2020).

Laboratory strategies to reduce AF aim to attenuate, remove, or chemically modify endogenous emitters (Wang et al., 2023). Common approaches include UV or white-light photobleaching (Neumann & Gabel, 2002), (Sapio et al., 2025), and chemical quenching with reagents such as sodium borohydride or Sudan Black B (Sun et al., 2017), (Wang et al., 2023). While effective in some contexts, these prospective procedures can be labour-intensive, may degrade tissue epitopes, and risk global signal loss, limiting their reproducibility across samples and platforms.

Beyond photochemical treatments, instrumentation hardware-assisted solutions have been developed. Fluorescence lifetime imaging microscopy (FLIM) can distinguish AF and TS based on differences in their fluorescence lifetimes (Becker, 2012), (Hwang et al., 2024), (Wang et al., 2025), while spectral imaging combined with linear unmixing (Zimmermann et al., 2003) exploits differences in emission spectra to decompose mixed signals. Although these approaches can achieve high precision, they are typically slower and less scalable than conventional widefield/confocal acquisition requiring specialized hardware modules, careful calibration, and advanced expertise.

Computational methods provide an attractive and broadly applicable alternative as they operate retrospectively, avoid additional wet-lab steps, and do not require instrument modifications. Existing approaches largely fall into subtraction-based and factorisation-based categories. Subtraction-based methods leverage unstained paired AF baselines that are subtracted from the mixed signal (MS) to estimate TS (Eng et al., 2022). These can be from the same channel in serial cycles, or from AF-dedicated channels in the same staining cycle. Interpolation-based AF subtraction (AFS) acquires baselines in the first and last cycles and estimates the corresponding AF of each cycle via linear interpolation (e.g., mplexable) (Eng et al., 2022), (Potier et al., 2022). These approaches rely on the assumption that AF decays linearly across imaging cycles and that this decay is spatially uniform within and across tissues, assumptions that as we later show in the manuscript, do not hold true in practice. More advanced subtraction methods include SAIBR, which regresses MS against an AF reference channel to estimate TS (Rodrigues et al., 2022), and AFid, which uses multichannel clustering on MS/AF pairs to identify and mask TS and AF regions (Baharlou et al., 2021). While effective in some settings, SAIBR presumes approximately linear correlations between AF and MS across channels, and AFid assumes that AF is shared across at least two channels and largely non-overlapping with TS. In tissues with multiple, spectrally distinct AF sources and complex staining patterns, these assumptions are frequently violated, as AF varies with tissue structure, fixation, and local biochemical composition. Factorization-based methods propose an alternative strategy. Non-negative matrix factorization (NMF) has been applied to decompose MS into AF and TS components, outperforming early linear unmixing methods (Woolfe et al., 2011). However, when AF dominates the variance, when AF contributions differ strongly between channels, or when AF and TS components are highly colinear, factorization-based approaches can misattribute AF by over-subtracting weakly expressed markers, under-subtracting channel-specific AF, or imprinting AF structure into the inferred TS component.

These challenges highlight the need for computational methods that can adapt to the different scenarios. To address this challenge, we present CLEAR-AF (Coordinate-transformed Local Estimation and Adaptive Removal of Autofluorescence), a novel subtraction-based framework that estimates AF at the pixel level. By transforming images into polar coordinates and applying Gaussian mixture modelling, CLEAR-AF captures spatial heterogeneity in AF distributions and adaptively subtracts background without sacrificing weak marker signals. We demonstrate that CLEAR-AF consistently improves AF removal across multiple tissue types and multiplexed imaging platforms, offering clearer molecular readouts for downstream single-cell analyses.

## Results

We evaluated AF removal on a multi-cancer tissue microarray (TMA) containing 30 formalin-fixed, paraffin-embedded (FFPE) cores from 30 human cancer samples: 13 melanoma (SKCM), 12 glioblastoma (GBM), 2 hepatocellular carcinoma (HCC), 2 breast cancer (BRCA) and one healthy tonsil (**Figure 1A**). Serial sections from this TMA were profiled using four cMIF platforms: MILAN (Bolognesi et al., 2017), COMET (LunaPhore), PhenoCycler Fusion (Akoya), and MACSima (Miltenyi). An overview of acquired protein markers and platform-specific acquisition parameters is provided in **Supplementary Table 1**. For MILAN, the same slide was scanned twice to assess the reproducibility of AF subtraction methods. The marker panel spanned a wide range of prevalence, from fully absent signal (no primary antibody, blank baselines) to highly prevalent staining (e.g., CD56 in GBM), thereby covering a broad spectrum of signal intensities and spatial patterns (**Figure 1B**).

**Figure 1.**
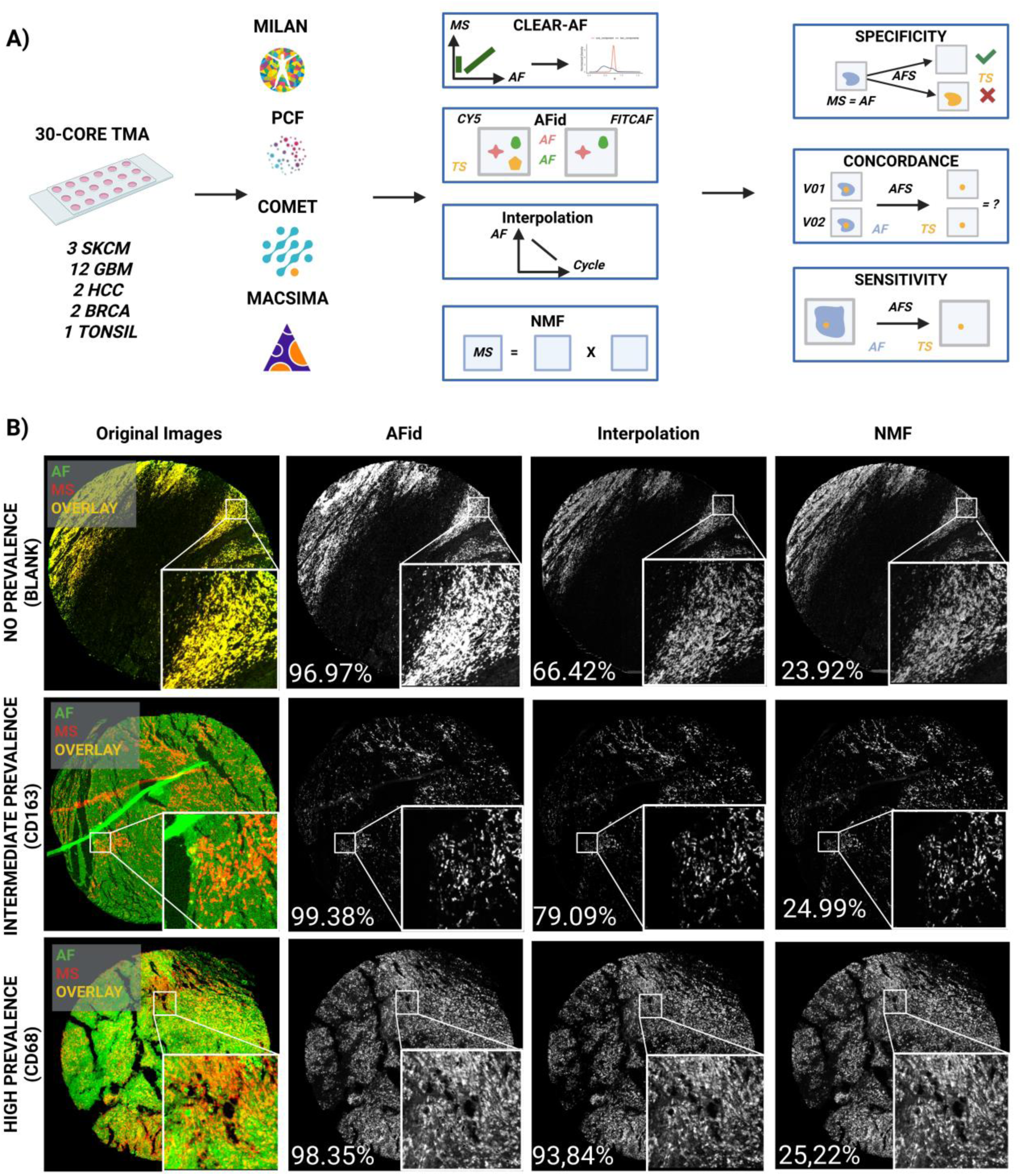
Overview of the experimental design and performance of existing autofluorescence subtraction (AFS) methods. Experimental design overview and limitations of the SOTA. A) A 30-core multi-cancer FFPE tissue microarray (TMA) comprising melanoma (SKCM), glioblastoma (GBM), hepatocellular carcinoma (HCC), breast cancer (BRCA), and tonsil tissue was serially sectioned and imaged using four cMIF platforms: MILAN, COMET, PCF, and MACSima. AFS was performed using three SOTA algorithms: AFid, interpolation-based subtraction, and NMF. Resulting images were evaluated in terms of specificity, sensitivity, and concordance. B) Representative examples illustrating three marker-prevalence scenarios: no prevalence (blank), intermediate prevalence (CD163), and high prevalence (CD68). For each case, original mixed-signal (MS) images and the corresponding AF estimates from AFid, interpolation, and NMF are shown. Percentages denote the proportion of estimated TS pixels over the baseline AF pixel count.

### Performance and variability of current AF removal methods

We first evaluated the performance of three state-of-the-art (SOTA) AFS methods: interpolation (Eng et al., 2022), (Potier et al., 2022), NMF (Woolfe et al., 2011), AFid (Baharlou et al., 2021) and evaluated their capacity to split MS images into AF and TS components. Visual inspection across blank, intermediate-prevalence (CD163), and high-prevalence (CD68) examples revealed substantial discrepancies between methods (**Figure 1B**). A quantitative assessment using Cohen’s kappa confirmed this observation: the agreement between methods was no better than chance, with mean pairwise k values of -0.004 for the no-prevalence example, -0.003 for the intermediate prevalence example, and 0.012 for the high-prevalence example. All the methods frequently underestimated the AF component, resulting in residual background in the TS channel. This under-subtraction was most pronounced in blanks and low-prevalence markers, where large areas of tissue introduced false positive (FP) signals, as reflected by the high fraction of pixels showing TS relative to the AF baselines (percentages reported in **Figure 1B**). Next, we assessed reproducibility using duplicate scans of the MILAN slide. Despite identical tissue, staining, acquisition, and image pre-processing, the three methods yielded noticeable different TS estimates between scans for the same core and marker (**Supplementary Figure S1**). Cohen’s kappa values further confirmed the lack of reproducibility, particularly in the subtraction-based approaches with k = 0.03 for interpolation, k = 0.009 for AFid, and k = 0.36 for NMF. This scan-to-scan variability suggests that current AFS approaches are sensitive to small acquisition differences and may not provide stable pixel-level estimates of TS. Together, these results indicate that existing AFS methods are neither sufficiently specific (especially in the absence of true staining), nor consistently reproducible across repeated acquisitions, highlighting the need for more robust AFS strategies.

### Development of CLEAR-AF

To address these limitations, we developed CLEAR-AF (Coordinate-transformed Local Estimation and Adaptive Removal of AutoFluorescence), a novel subtraction-based method that adaptively estimates the AF contribution at the pixel level using a geometry-driven framework (see methods). Briefly, each pixel in the images is represented in a two-dimensional Euclidean space defined by its AF and MS fluorescence intensities. This space is then transformed into polar coordinates (ρ, φ) (**Figure 2A**). The resulting φ image encodes, for each pixel, the angle between AF and MS; larger φ values correspond to pixels where MS is relatively enriched compared with AF and are therefore more likely to contain TS (**Figure 2B**).

**Figure 2.**
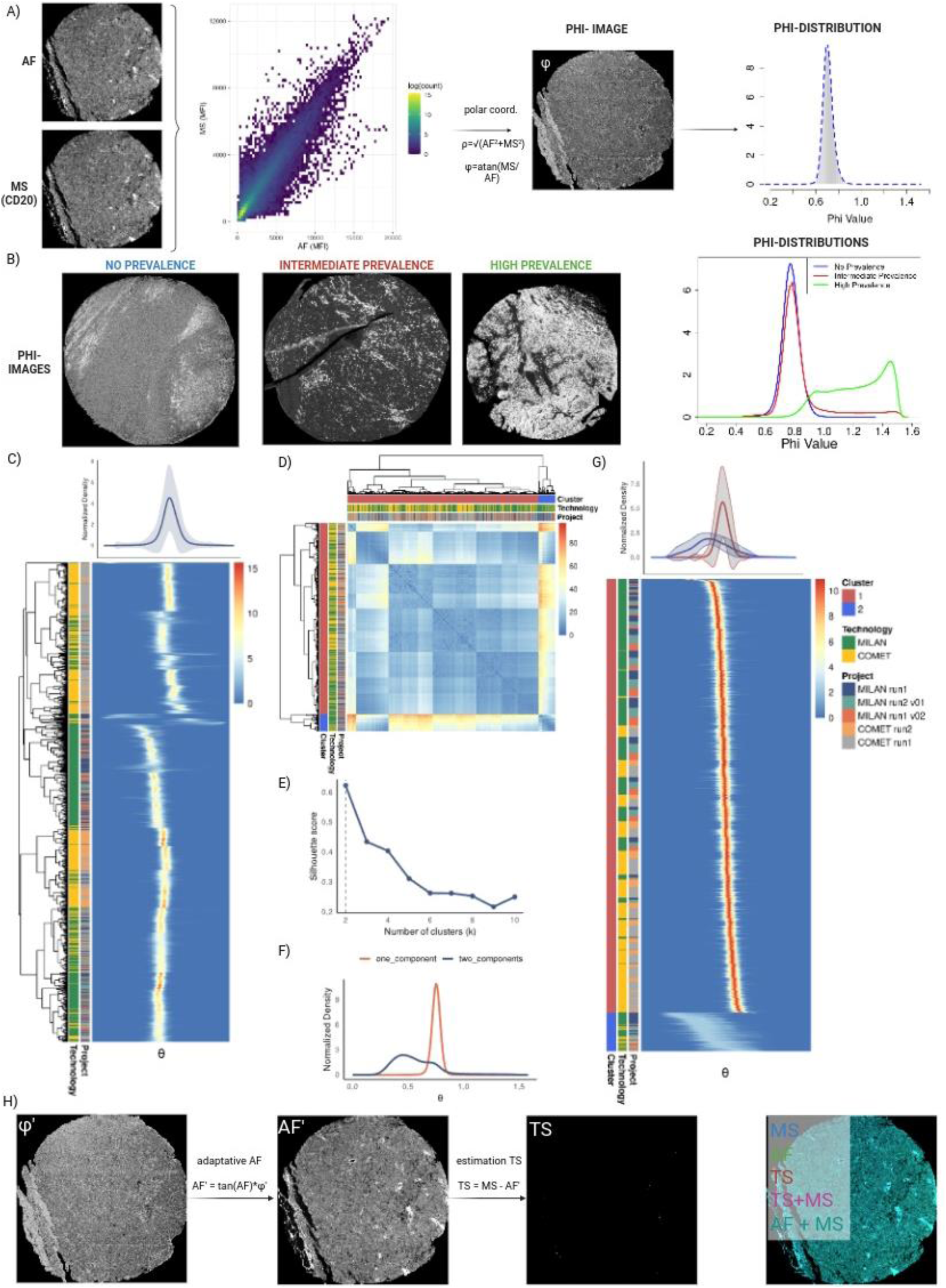
Geometry-driven modelling of φ distributions and adaptive AF subtraction in CLEAR-AF. **A)** Schematic of the coordinate transform. For each pixel, AF and MS intensities define a point in Euclidean space that is mapped to polar coordinates (ρ, φ). The φ image encodes the angle between AF and MS for every pixel. **B)** Representative φ images (left) and corresponding φ distributions (right) for the three cases shown in Figure 1B. **C)** Heatmap of smoother, normalised φ histograms across 1,290 MILAN and COMET images, ordered by hierarchical clustering. **D)** Pairwise dynamic time warping (DTW) distance matrix between φ histograms with hierarchical clustering. **E)** Silhouette analysis indicating an optimal cluster number of k=2. **F)** Density plots of the two cluster medoids, used as reference templates for subsequent classification. **G)** All φ distributions aligned to the medoid of their assigned cluster, confirming two distinct archetypical shapes: Class 1 (narrow and unimodal), and Class 2 (broad and multimodal). **H)** Schematic of adaptive AF subtraction. The φ image is trimmed according to the class-specific threshold to obtain φ’, which is used to reconstruct the AF component of the MS image (AF’). TS is then estimated as TS = MS – AF’. Example images illustrate, φ’, AF’, TS, and composite overlays of MS, AF and TS channels for the same case depicted in section A.

The distribution of φ values corresponding to the three representative cases in **Figure 1B** illustrates how signal prevalence shapes this angular representation (**Figure 2B**). In the absence of TS, φ values follow a narrow unimodal distribution. When TS is rare, the distribution remains approximately unimodal but exhibits a skewed tail towards higher φ values. In contrast, images with high TS prevalence typically display a broader, often bimodal, φ distribution. This observation motivates using the φ distribution itself to guide AF subtraction.

To systematically characterise this landscape, we analysed φ distributions for 1,290 MILAN and COMET images spanning multiple tissues and markers (**Figure 2C**). Because absolute φ locations vary substantially across images, for example, due to changes in AF/MS ratios across cycles (see Dynamics of tissue autofluorescence in cyclic multiplexing experiments), we focused on the shape of the distributions rather than in their absolute position. We computed pairwise dynamic time warping (DTW) distances between smoothed φ histograms and performed hierarchical clustering on the resulting distance matrix (**Figure 2D**). Silhouette analysis identified two as the optimal number of clusters (**Figure 2E**). The corresponding cluster medoids, which serve as reference templates for the classification of new query images, are shown in **Figure 2F**. Aligning individual φ distributions to their respective medoid revealed two clear archetypes (**Figure 2G**). Class 1 distributions are narrow and unimodal, characteristic of images without or with low prevalence of TS. Class 2 distributions are broader and frequently bimodal, characteristic of images with high prevalence of TS. CLEAR-AF employs distinct thresholding strategies for these two classes. For Class 1, φ is modelled as a single Gaussian, and the AF-TS threshold is defined as two standard deviations above the mean. For Class 2, φ is modelled as a two-component Gaussian mixture, and the threshold is set at the intersection of the two components (see methods). This threshold is then used to truncate the φ image, yielding a trimmed angle map (φ’) from which the AF component of the MS image (AF’) is reconstructed, Finally, TS is estimated on a per-pixel basis as the difference between MS and AF’ (**Figure 2H**).

### Benchmarking CLEAR-AF against SOTA AFS methods

We benchmarked CLEAR-AF against three SOTA AFS methods (interpolation (Eng et al., 2022), (Potier et al., 2022), NMF (Woolfe et al., 2011), AFid (Baharlou et al., 2021)) using two criteria: (i) concordance between repeated stainings on the same MILAN slide, and (ii) specificity (under-subtraction, e.g., resilience to false positives, estimating TS signals where there should be AF).

#### Evaluation of concordance

To assess scan-to-scan consistency, we compared TS maps obtained from two independent MILAN acquisitions (v01 and v02) on the same slide. This was quantified using Cohen’s kappa which measures the level of agreement between two observations (1 is perfect agreement, 0 is chance-level agreement, -1 is perfect disagreement). CLEAR-AF produced substantially more consistent TS estimates (mean kappa = 0.36) across all v01/v02 scans, followed by interpolation (mean kappa = 0.25, p-adj < 0.0001), AFid (mean kappa = 0.13, p-adj < 0.0001) and NMF (mean kappa = 0.026, p-adj<0.0001) (**Figure 3A, C**).

**Figure 3.**
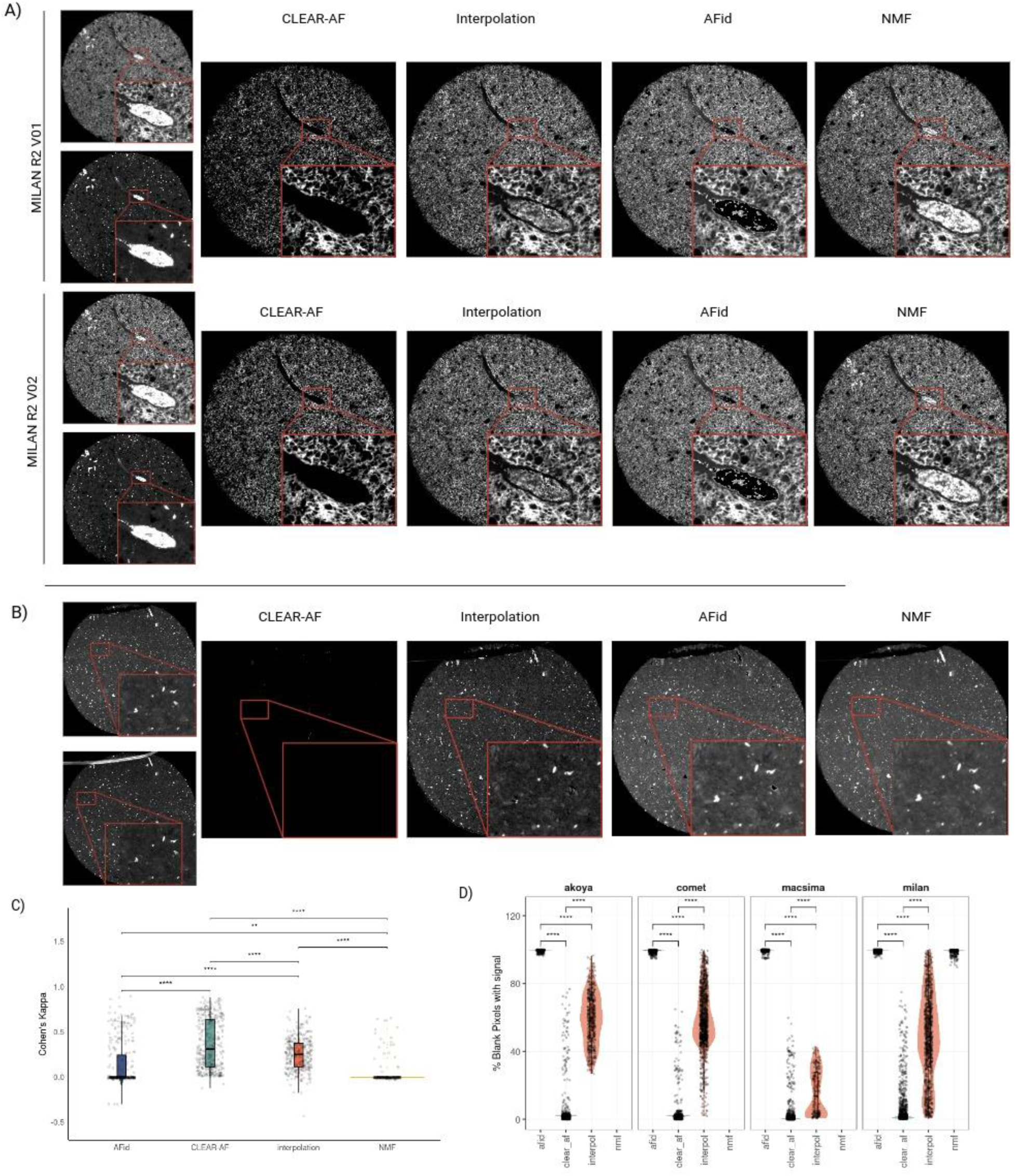
CLEAR-AF improves concordance and specificity compared with existing AFS methods. **A)** Representative MILAN core illustrating TS estimates obtained with CLEAR-AF, interpolation, and AFid for two independent scans of the same slide (v01 and v02). **B)** Representative blank core showing AF-only images (left) and the corresponding TS estimates after AFS with CLEAR-AF, interpolation, and AFid. CLEAR-AF largely removes background, whereas interpolation and AFid leave substantial residual signal. **D)** Violin plots of Cohen’s κ statistics quantifying concordance between MILAN v01 and v02 for each AFS method. Each point corresponds to a marker–core pair. **D)** Violin plots showing the percentage of pixels classified as TS in blank images across all non-DAPI channels, stratified by imaging platform (Akoya, COMET, MACSima, MILAN). Lower values indicate better specificity (fewer false positives). Statistical significance was assessed using Wilcoxon rank-sum tests with FDR correction (*p < 0.05; **p < 0.01; ***p < 0.001; ****p < 0.0001).

#### Evaluation of specificity

We next evaluated specificity using blank images, where no TS is expected and any residual signal after AFS reflects false positives. For each dataset, we quantified the fraction of pixels classified as TS in non-DAPI channels (**Figure 3B, D**). CLEAR-AF consistently yielded the lowest false-positive rates (AKOYA = 4.02%, COMET = 2.36%, MACSIMA = 1.38%, MILAN = 4.07%), markedly lower than interpolation (AKOYA = 59.00%, COMET = 58.74%, MACSIMA = 15.87%, MILAN = 47.88%, p-adj < 0.0001 in all cases), AFid (AKOYA = 99.39%, COMET = 99.16%, MACSIMA = 99.48%, MILAN = 99.44%, p-adj < 0.0001 in all cases) and NMF (MILAN = 98.8%, p-adj < 0.0001).

Together, these analyses show that CLEAR-AF achieves higher concordance across repeated acquisitions and markedly better specificity on blanks than existing AFS methods, indicating more stable and reliable estimation of TS.

In our experiments, more than one AF baseline was available for a given marker. We observed that the choice of baseline can substantially affect the resulting TS estimate. **Figure 4** illustrates this effect for a FOXP3 staining (cycle 5, R05) in the TRITC channel, where two AF references (R01 and R06) were used for AFS. In these cases, CLEAR-AF produces markedly different φ distributions (**Supplementary Figure S2**). The φ histogram obtained with R06 is narrower than with R01 (coefficient of variation: *CV*_*R*01_ = 7.37, *CV*_*R*06_ = 2.90), indicating a more homogeneous and stable AF–to–MS relationship across the tissue. Using R06 as the reference therefore yields a cleaner TS estimate with fewer spurious signals (*TS*_*R*01_ = 0.59% of foreground pixels vs. *TS*_*R*06_ = 0.25%; **Figure 4**). Visual inspection of the vessel region confirms that the AF pattern has changed across cycles affecting the subtraction.

**Figure 4.**
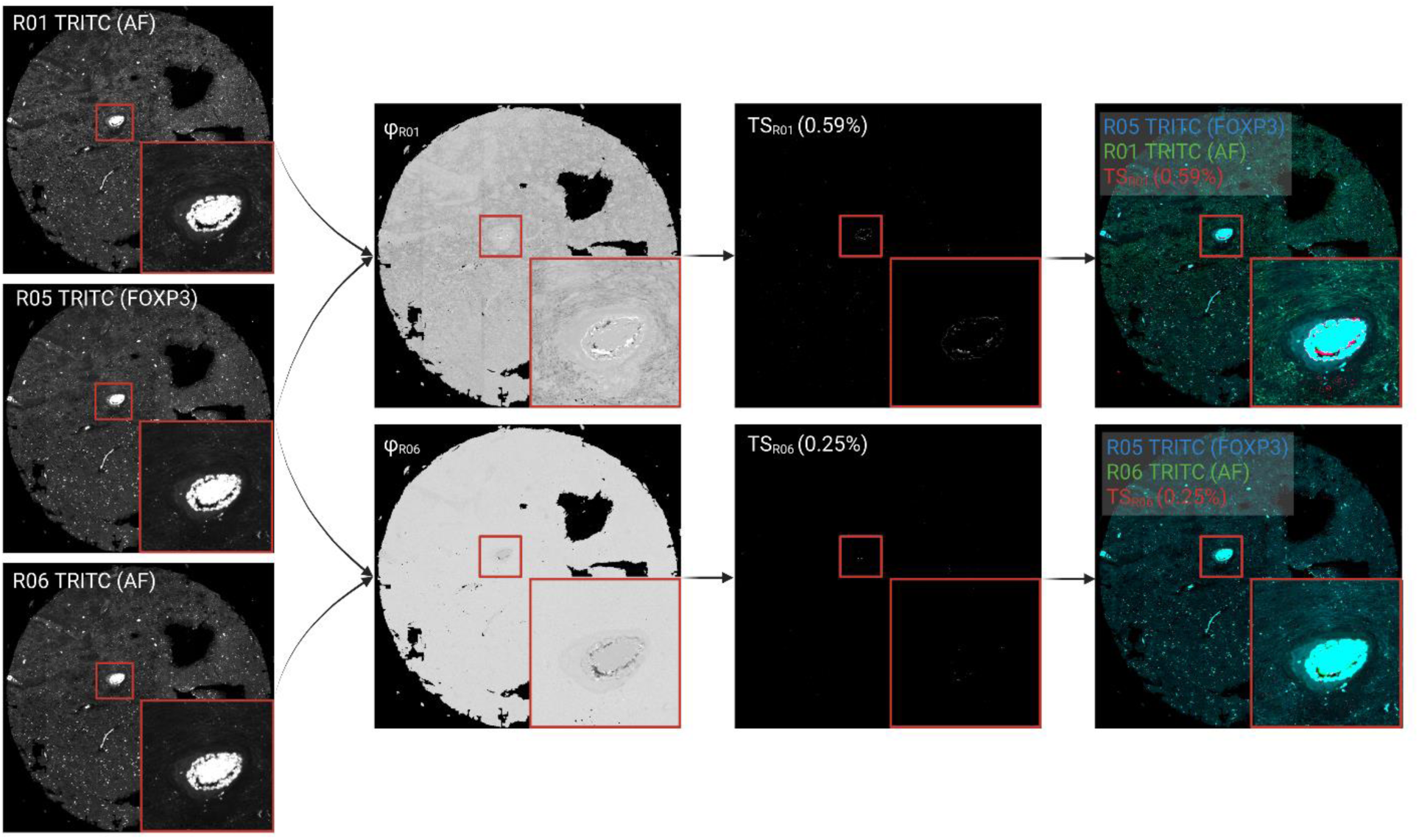
Effect of AF baseline selection on CLEAR-AF subtraction. Left column: AF reference images acquired in the TRITC channel (R01 and R06) and the corresponding FOXP3 staining (R05, TRITC) used as mixed signal (MS). Zoomed areas highlight a perivascular region. Centre-left column: φ images computed by CLEAR-AF when pairing the FOXP3 MS image with AF baselines R01 (top) or R06 (bottom). Centre-right column: the resulting TS estimates. Percentages indicate the fraction of foreground pixels with TS. Right column: overlays of MS (cyan), AF baseline (green), and TS (red) for the two AF choices. Using R06 as baseline leads to a narrower φ distribution, fewer TS pixels, and visually cleaner subtraction in the perivascular region than using R01.

### Dynamics of tissue autofluorescence in cyclic multiplexing experiments

In this light, we ought to investigate the dynamics of tissue autofluorescence in cMIF. cMIF enables comprehensive spatial profiling by iteratively labelling and imaging a tissue sample across multiple staining cycles. However, repeated exposure to excitation light and wet-lab processing steps can compromise signal fidelity over time, leading to cumulative photobleaching, tissue degradation, and variable intensity decay across markers, cycles, and spatial regions (Demchenko, 2020).

To assess these effects, we systematically analyzed signal stability across cycles for all datasets, quantifying both global and regional changes in median intensity. **Figure 5** illustrates these dynamics for the COMET slide in the DAPI channel. Across the 24 scan regions, median intensity decreased by ∼20% on average over 18 cycles, but with substantial variability between samples (-4.82% for S9 to -60% for S23). Importantly, this progressive signal loss is not linear with each tissue sample following a different dynamic trajectory (**Figure 5A-C**). Summarising the cross-cycle variability across all platforms and acquisition channels, MACSIMA exhibited the most stable performance (mean 0.12% intensity change per cycle), followed by COMET (0.45%), AKOYA (1.32%), and MILAN (2.23%) (**Figure 5D, E**). These differences indicate that AF and TS dynamics are tissue and platform dependent and cannot be captured by a single uniform decay model.

**Figure 5.**
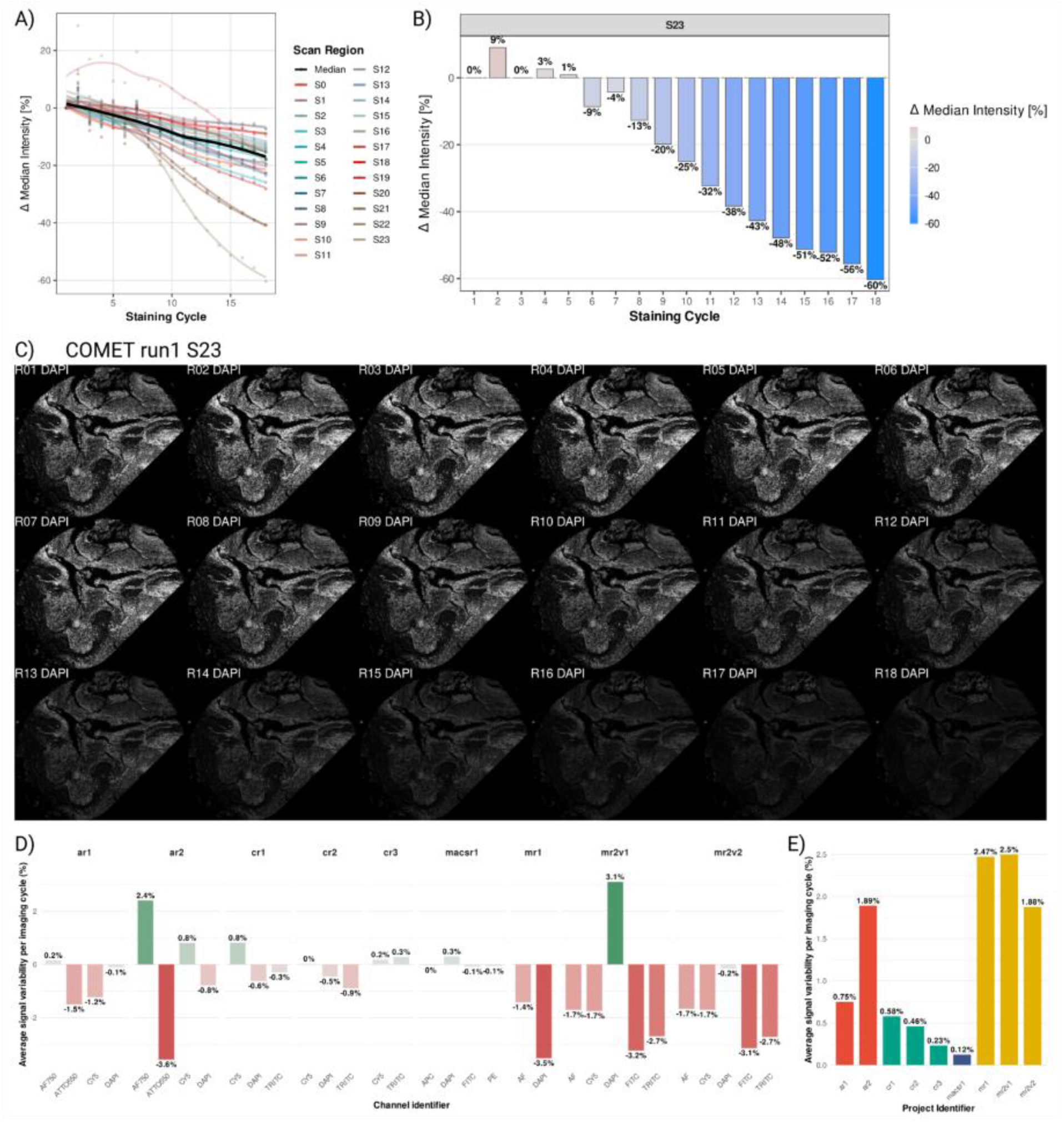
Cross-cycle signal stability across scan regions and acquisition technologies. **A)** Median DAPI intensity change across staining cycles for 24 samples (scan region) from COMET. Each line represents a sample; the black line denotes the median. **B)** Per-cycle median DAPI intensity for scan region S23, illustrating a 60% decrease over 18 cycles. **C)** Raw DAPI images for S23 from cycles R01–R18, displayed with a shared intensity window to visualize progressive signal loss. **D)** Mean per-cycle signal variability for each acquisition channel and technology, expressed as percentage change in median intensity. Color shading encodes variability of magnitude. **E)** Mean per-cycle signal variability aggregated by acquisition technology. Bar colors indicate acquisition platform (red: AKOYA, green: COMET, blue: MACSIMA, yellow: MILAN), showing platform-specific differences in signal stability.

To further dissect intra-tissue behaviour, we evaluated spatial AF dynamics in a tissue- and region-specific manner. Using the baseline AF image (cycle 1), we applied k-means clustering (k = 10) to segment the section into regions with similar initial AF intensity and then tracked the median φ value for each cluster across cycles. **Supplementary Figure 3** shows an example from COMET run2, S11 (Cy5 channel). Tumour-rich clusters (K09–K10) exhibited increasing signal over time (+10.2% and +19.8%), whereas stromal and vascular clusters (K01–K08) showed progressive loss, with an average decrease of –7.4%.

Together, these analyses demonstrate that AF dynamics vary not only between platforms and tissues, but also across regions within a single section. AF decay is therefore not spatially uniform. Assuming a global, homogeneous decay or applying a single scaling factor across an entire slide risks introducing systematic bias in signal estimation, particularly in long cMIF experiments with heterogeneous tissue architecture. These findings support the need for tissue-aware AF modelling and for selecting AF baselines that are regionally representative when calibrating and normalising subtraction methods such as CLEAR-AF.

## Conclusion

cMIF enables the acquisition of tens to hundreds of protein markers at single-cell resolution, substantially expanding our understanding of spatial biology in recent years (Karimi, 2024). However, these imaging data are highly susceptible to background signals from intrinsically fluorescent tissue components (including collagen, elastin, NAD(P)H, and lipofuscin) that emit fluorescent signals upon excitation at specific wavelengths and spectrally overlap with fluorophores targeting proteins of interest (Monici, 2005), (Lakowicz, 2006), (Richardson & Lichtman, 2015). Accurate separation of protein-target and autofluorescent signals is therefore essential for reliable downstream analyses (Chevrier et al., 2018).

In this study, we systematically examined the behaviour of autofluorescence across tissues, platforms, regions, and staining cycles, showing that AF dynamics are neither uniform nor predictable using simple global models. Existing computational approaches frequently fall short in terms of sensitivity, specificity, and/or concordance, highlighting the need for novel strategies.

To overcome these challenges, we introduced CLEAR-AF, a geometry-driven framework that adaptively estimates and subtracts AF on a per-pixel basis. By transforming AF and MS intensities into polar coordinates and modelling the resulting φ distributions, CLEAR-AF captures the underlying AF-MS-TS structure and automatically adjusts to the prevalence and spatial heterogeneity of true signal. This adaptive design enables robust performance across a broad range of markers, tissues, and imaging platforms, outperforming SOTA methods in terms of sensitivity, specificity, and concordance.

In brief, CLEAR-AF establishes a new, platform-agnostic solution for reliable autofluorescence subtraction in cyclic multiplexed imaging, advancing the field towards more reproducible and trustworthy spatial proteomics.

## Materials and Methods

### Dataset description

This study used a tissue microarray (TMA) block containing 30 formalin-fixed, paraffin-embedded (FFPE) cores from 30 different human samples: 13 melanoma samples (SKCM), 12 glioblastoma (GBM), 2 hepatocellular carcinoma (HCC), 2 breast cancer (BRCA), and one healthy tonsil. Serial sections from this TMA block were cut and imaged using four different cMIF technologies: the MILAN protocol (Bolognesi et al., 2017), COMET (Lunaphore Technologies), Phenocycler Fusion (Akoya Biosciences) and Macsima (Miltenyi Biotec) (**Figure 1A**). For the MILAN dataset, the same stained slide was scanned twice to assess the reproducibility of the different AFS methods. The experimental design for each acquisition technology, including the number of cycles, acquired markers, acquisition channels, and technical details regarding each channel (excitation/emission wavelengths, exposure times) are detailed in **Supplementary Table 1**.

### Image pre-processing

All images underwent the same pre-processing pipeline prior to the application of any AFS method. First, flat-field correction was performed using the BaSiC algorithm (Peng et al., 2017), applied independently to each cycle and fluorescence channel. Second, stitching and registration were performed using COLLAGE (Antoranz et al., 2024), resulting in whole-slide image stacks in which all cycles were spatially aligned. From these registered whole-slide images, individual TMA cores were extracted using a custom de-arraying pipeline that identified sample regions. The resulting core-level, cycle-aligned image stacks constituted the input data for all autofluorescence subtraction methods evaluated in this study.

### CLEAR-AF Method

In fluorescence imaging, images acquired with antibodies targeting specific proteins (measured signal, MS) acquired fluorescence from both the protein of interest (true signal, TS), as well as from intrinsically fluorescent components present in the tissue (autofluorescence, AF). Therefore, MS at a given pixel can be expressed as a combination of AF and TS contributions,

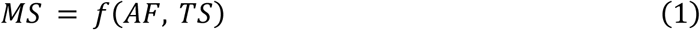

Where MS is the measured image, AF is the autofluorescent component of MS, and TS is the true signal component of MS. In cMIF, subtraction-based approaches use baseline images acquired without primary antibodies which include only AF signals (AF’). These proxies are used to estimate TS (TS’) by subtracting AF’ from MS,

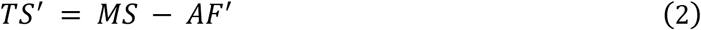

Where AF’ is the fluorescence acquired in the baseline image, and TS’ is the true signal estimated by subtraction. However, as shown in the manuscript, these baseline images do not correspond 1-to-1 to the true AF in MS given the AF degradation throughout the cycles on top of stochastic differences in acquisition (Monici, 2005), acting as a proxy. Additionally, as shown in the manuscript, this degradation is not linear. Therefore, direct or scaled interpolation-based subtraction is not specific enough for accurate TS estimation. Consequently, in CLEAR-AF (Coordinate-transformed Local Estimation and Adaptive Removal of AutoFluorescence), we first project the co-registered AF’, MS fluorescence values for each pixel to a Euclidean space. A textbook case of a scatterplot of these values reveals two main distributions: one in which AF’ and MS intensities are strongly correlated (representing AF components of MS), and one where MS intensity is largely independent of AF (representing pixels MS pixels with both AF and TS) (**Figure 6**). The goal is therefore to identify these two components and use them to model the AF component of MS in the second category. To that end, we transform this Euclidean space into polar coordinates,

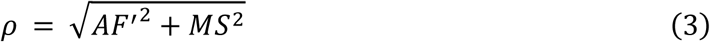

**Figure 6.**
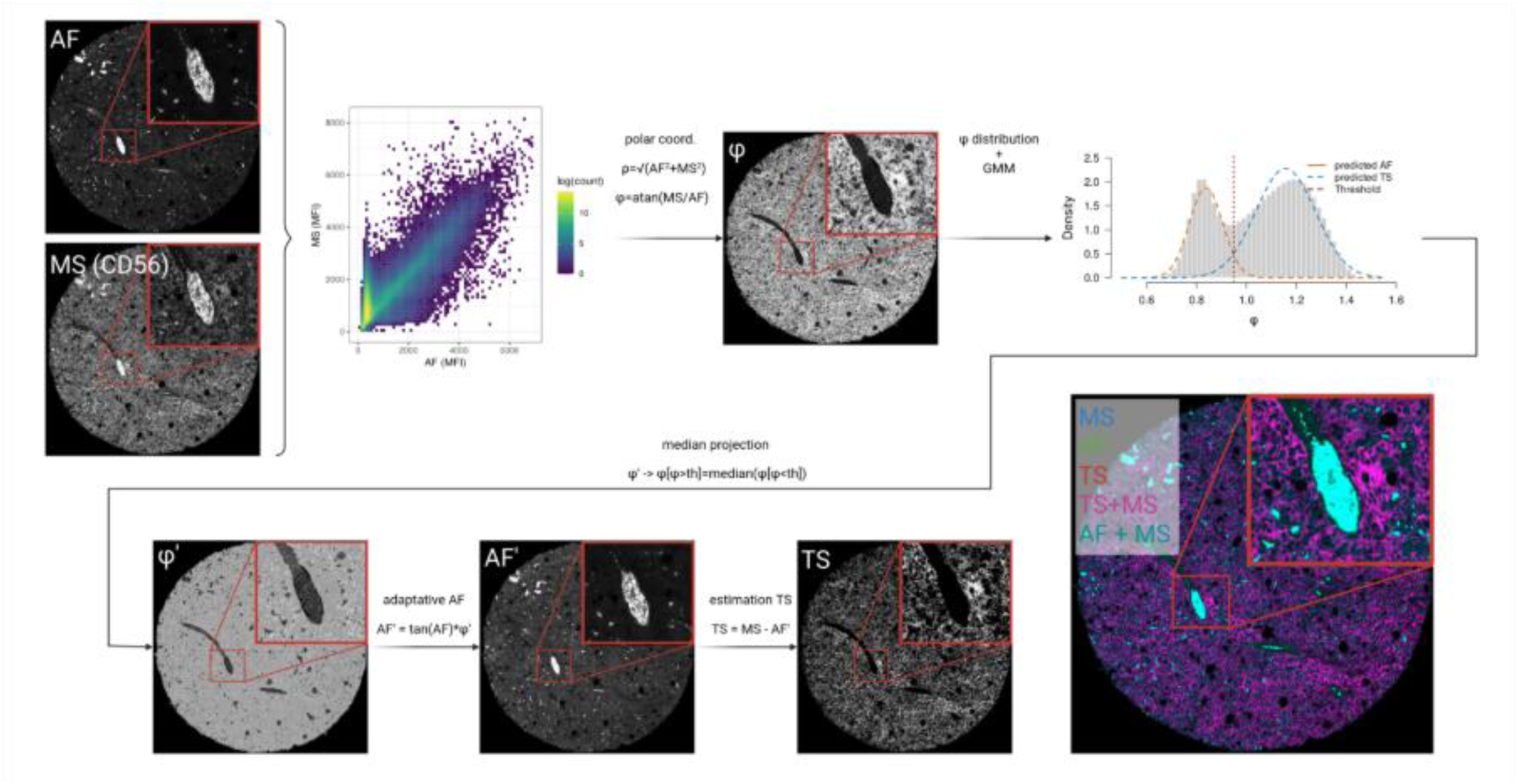
CLEAR-AF: geometry-aware subtraction of autofluorescence in multiplexed imaging. Overview of the CLEAR-AF pipeline for estimating true signal (TS) from mixed signal (MS) and autofluorescence (AF). Left: Input images consist of an AF-only image and a MS image for the same sample and registered (for MS, CD56 marker in a GBM sample is shown). Middle: A joint intensity scatter plot (AF vs MS) is transformed into polar coordinates, and the φ distribution is modelled using Gaussian mixture models (GMM) with 1 or 2 components to separate AF- and TS-components. Lower branch: Pixels above the GMM- or Gaussian-derived threshold are projected onto the AF manifold via the median projection. The projected φ (φ‘) is used to estimate the AF component of MS (AF’), and TS is obtained by subtracting it from MS (TS’ = MS – AF’) Right: Composite image displaying MS (blue), AF (green), and TS (red), illustrating accurate recovery of CD56 signal.

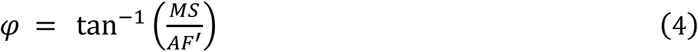

where, ρ denotes the radial intensity and φ encodes the angle between AF’ and MS. A higher φ implies greater likelihood that the pixel contains TS. For a given image, we model the empirical distribution of φ using a Gaussian Mixture Model (GMM) with two components, corresponding to AF-dominated and TS-dominated pixel populations. The threshold *φ*_*th*_ separating these two components is defined as the intersection point of the fitted Gaussians. Pixels with *φ* ≤ *φ*_*th*_ are considered AF, while pixels with *φ* > *φ*_*th*_ are considered to have both AF and TS signals. To split them, the corresponding φ value of those pixels is projected back onto the AF distribution by replacing it with the median value of the AF distribution. The projected φ value (φ’) is therefore:

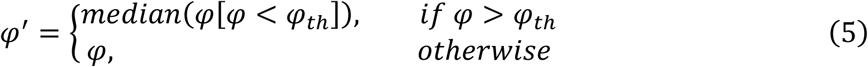

Using this projected angle, we compute the adaptive AF contribution per pixel as:

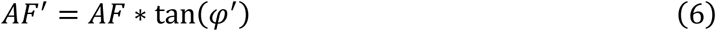

Substituting this on equation (2), allows us to obtain TS’. While this approach can effectively model cases with high signal prevalence where φ shows a clear bimodal distribution (**Figure 6**), it leads to under-subtraction in cases with no/intermediate prevalence where φ is clearly unimodal (**Supplementary Figure 4**). In these cases, we instead model the φ distribution as a single Gaussian, and define the cutoff at a one-sided 95% confidence level:

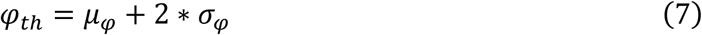

Where *μ*_*φ*_ and *σ*_*φ*_ are the mean and standard deviation of the φ distribution. To assess the generalizability of this approach, we analyzed the φ distribution across 1,290 AF/MS image pairs acquired with the MILAN and COMET technologies. In each case, the density function of the φ distribution was smoothed using a rolling mean (window size = 5) to reduce noise while preserving overall shape, and renormalized so that the area under the curve equals 1. As the specific ratio between AF and MS can vary significantly across cycles (**Supplementary Figure S2**), we aimed to identify common shapes in φ distributions regardless of their positional shifts. To that end, we employed dynamic time warping (DTW) as a shape-aware distance metric, which enables the alignment of density functions with locally shifted features, allowing comparison based on overall shape rather than absolute position. This approach effectively accounts for the large shifts observed across φ distributions. Pairwise DTW distances between smoothed density functions were computed using the *dtwclust* R package (Giorgino, 2009), providing a shape-sensitive distance matrix for subsequent clustering analyses. To limit over-alignment across unrelated regions and improve computational efficiency, a constrained warping window (*Sakoe-Chiba band, w=5)* was applied when computing the DTW distances. Hierarchical clustering was then applied on the DTW distance matrix with varying number of clusters (from 2 to 10). The optimal number of clusters was selected using the Silhouette Index. For each cluster, the medoid (the density with minimal average DTW distance to all other members) was taken as a representative templated and used to assign new images to the appropriate class (bimodal vs unimodal), thereby selecting the corresponding CLEAR-AF thresholding strategy. An overview of CLEAR-AF is provided in **Figure 6**.

### Benchmarking against the SOTA

CLEAR-AF was benchmarked against three widely used autofluorescence removal approaches: interpolation-based subtraction (Eng et al., 2022), (Potier et al., 2022) but also implemented in the Horizon (LunaPhore) and InForm (Akoya) softwares; NMF based autofluorescence removal (Woolfe et al., 2011); and AFid (Baharlou et al., 2021).

### Interpolation-based subtraction

The interpolation-based method is a weighted interpolation between an initial and final baseline cycle. In cMIF experiments, the signal intensity can degrade over successive cycles due to photobleaching, repeated exposure to excitation light, tissue degradation, and other wet-lab related factors (Demchenko, 2020). Therefore, many subtraction-based methods compensate for this variability by scaling the AF image with a factor α, applied globally to all pixels:

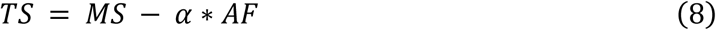

Where α is the scaling factor and calculated by linearly interpolating the intensity values of the baseline images acquired at the beginning (first cycle, *AF*_1_) and at the end (last cycle, *AF*_*n*_) of a cMIF experiment:

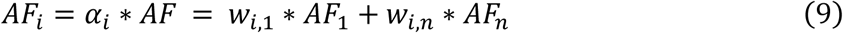

Where *w*_*i*,1_ and *w*_*i*,*n*_ are the weights assigned to the two baselines:

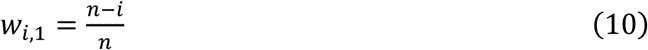

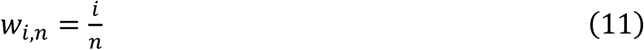

This approach was implemented using a custom script in Python.

### AFid

AFid identifies autofluorescent structures using information from two fluorescence channels. First, each image is smoothed with a Gaussian filter and thresholded with a Niblack local threshold, producing two binary masks. Their intersection yields an “intersection mask” containing candidate fluorescent objects. Features such as correlation and kurtosis are then extracted for each object, standardized, and clustered using *k-means*. The cluster with the highest correlation is designated as AF. Pixels assigned to AF are removed from the images. AFid was implemented following the AFid R package.

### NMF-based autofluorescence removal

In addition to subtraction-based strategies, we implemented the factorization method proposed by Woolfe et al. (Woolfe et al., 2011), which uses non-negative matrix factorization (NMF) to model MS as a combination of TS and AF with additive and multiplicative factors.

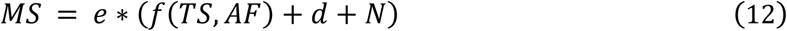

Where e is the exposure time, d is the dark current offset as an additive factor and N the noise. The function f(TS, AF) is modelled as a linear mixture:

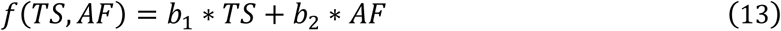

Where *b*_1_and *b*_2_are coefficients expressing the contribution of TS and AF to MS, respectively. In a matrix notation this can be rewritten as follows:

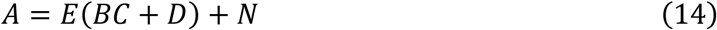

Where A contains the observed images, E is the diagonal matrix of exposure times, B are the mixture weights, C=[TS;AF] contains the latent TS and AF components, D contains the dark-current offsets, and N is Gaussian noise. The goal of the method is to estimate B, C, and D from the observations A and known exposures E. Please note that here A is a multi-channel image across different acquisition channels. A custom pipeline implementing this method was developed in Python.

### Evaluation strategies

All AFS methods were benchmarked under three complementary evaluation scenarios: (i) concordance between repeated acquisitions of the same slide, and (ii) specificity on baseline (AF-only) images.

### Evaluation of concordance

To quantify how consistently each method estimates TS across repeated acquisitions, we computed Cohen’s kappa (k) between AFS-derived TS masks from two independent scans of the same slide acquired with the MILAN platform (v01 and v02). Cohen’s kappa measures the agreement between two observations after accounting for the chance that they could agree randomly, providing a more robust measure than simple percent agreement, with a value of 1 indicating perfect agreement, 0 indicating chance-level agreement, and negative values suggesting worse-than-chance agreement. For each marker and TMA core, we first binarized TS images (TS > 0) to separate between pixels where the AFS method identified TS and pixels where the AFS method only identified AF. We limited this evaluation to the foreground areas where there is tissue present.

For every pair of TS masks, we constructed a confusion matrix at the pixel level where: true positives (TP) are defined as pixels classified as TS in both versions (intersection of both TS masks); false positives (FP) are defined as pixels classified as TS in v01 but not in v02; false negatives (FN) are defined as pixels classified as TS in v02 but not in v01; and true negatives (TN) are defined as pixels within the tissue mask that are classified as AF in both versions. Cohen’s k is then computed as:

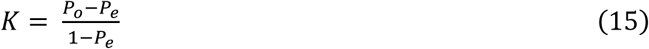

Where *P*_*o*_ is the observed agreement, and *P*_*e*_ is the expected agreement by chance and can be calculated as:

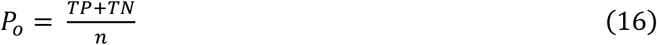

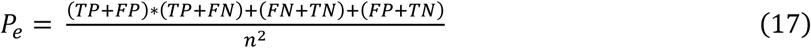

where *n* = *TP* + *FP* + *FN* + *TN*.

Here, kappa was computed using the kappa function in the vcd R package (default unweighted settings).

### Evaluation of specificity

To assess the ability of each method to avoid spurious signal, we evaluated specificity on AF-only (blank baseline) images, i.e., cycles/channels acquired without primary antibodies. For each blank image, we applied the different AFS methods and computed the proportion of tissue pixels that were still having signal after subtraction (TS>0). Because blank images contain no true specific staining by design, any residual TS pixels represent false positives. Formally, for each blank image we calculated:

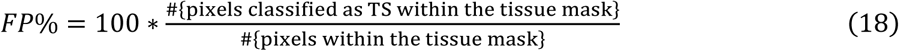

Lower FP% indicates better specificity.

---

**Supplementary Figure S1.**
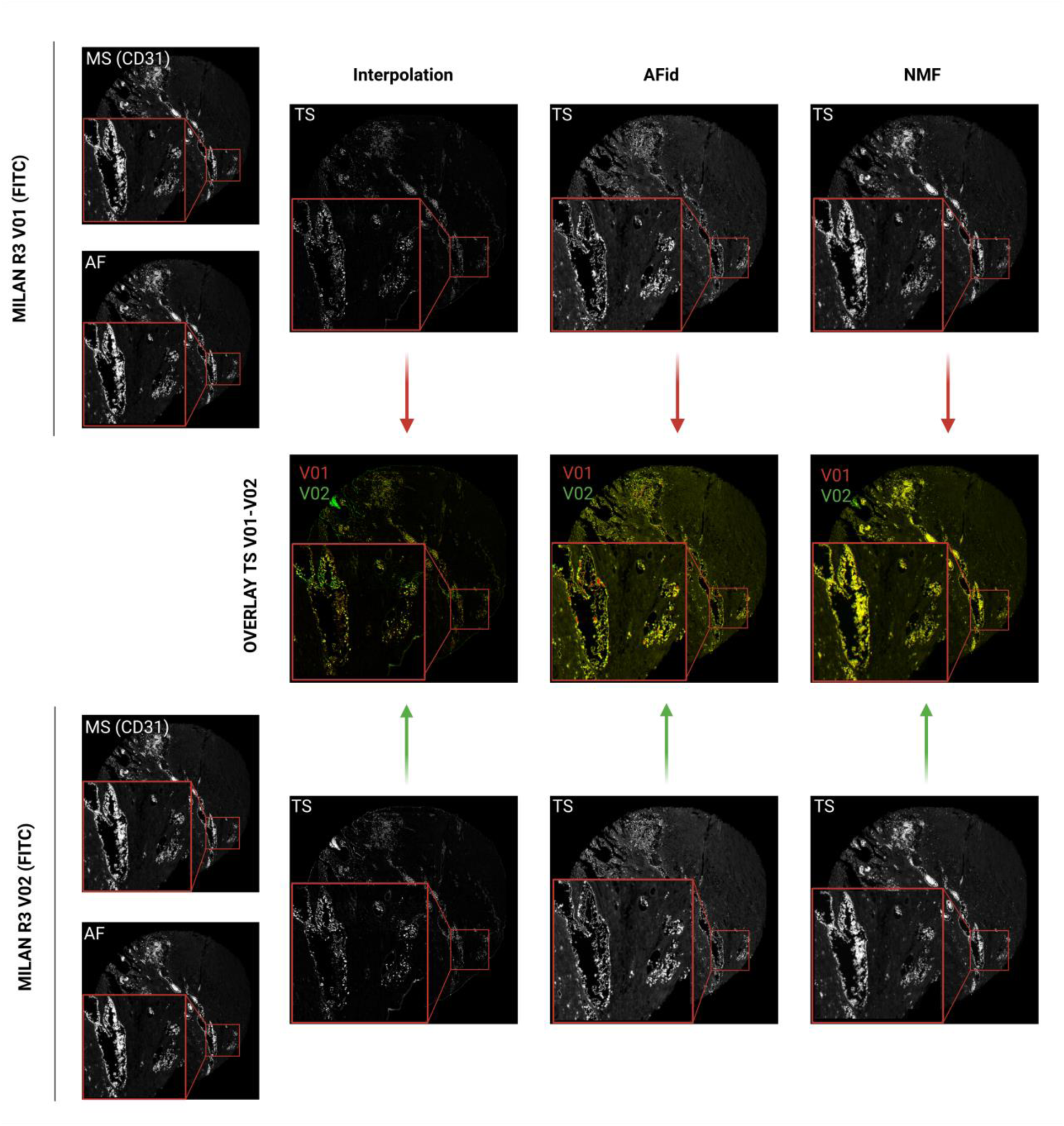
Reproducibility of AFS methods across duplicate scans of the same slide. Example illustrating the variability in true-signal (TS) estimates produced by interpolation, AFid, and NMF when applied to two independent scans (v01 and v02) of the same MILAN slide and core (marker CD31, FITC). Top row: mixed-signal (MS) and autofluorescence (AF) inputs for v01 and corresponding TS outputs for each AFS method. Bottom row: MS, AF, and TS outputs for v02. Middle row: pixel-wise overlays of TS masks from v01 (red) and v02 (green).

**Supplementary Figure S2.**
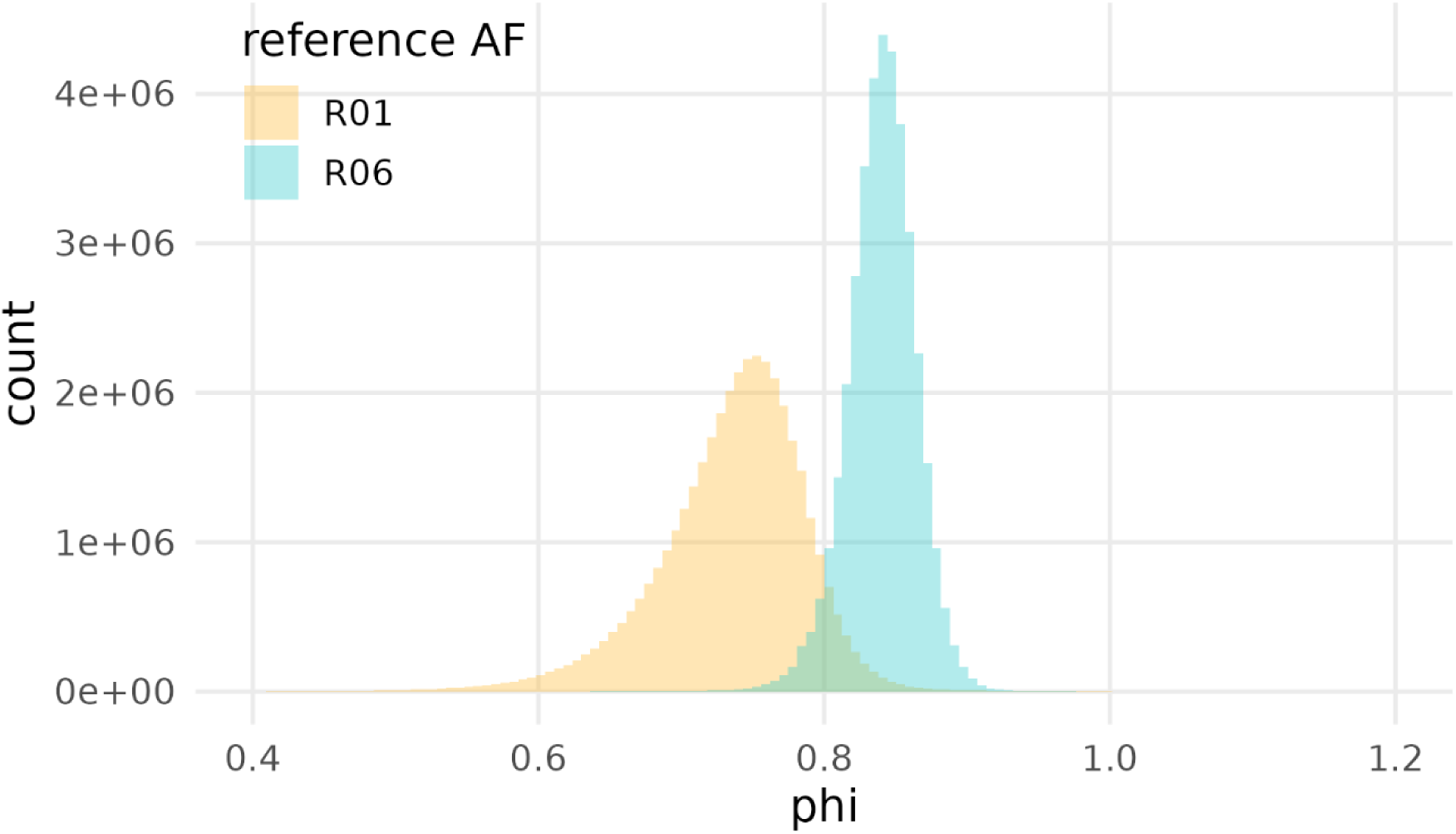
φ distributions for alternative AF baselines. Histogram of φ distributions obtained when pairing the FOXP3 MS image with AF baselines R01 (yellow) and R06 (cyan). The R06-based φ distribution is narrower (CV_φ_r01) = 7.37 vs. CV_φ_r06 = 2.90), indicating a more consistent AF–to–MS relationship and motivating the choice of R06 as the preferred baseline for CLEAR-AF subtraction in Figure 4.

**Supplementary Figure 3.**
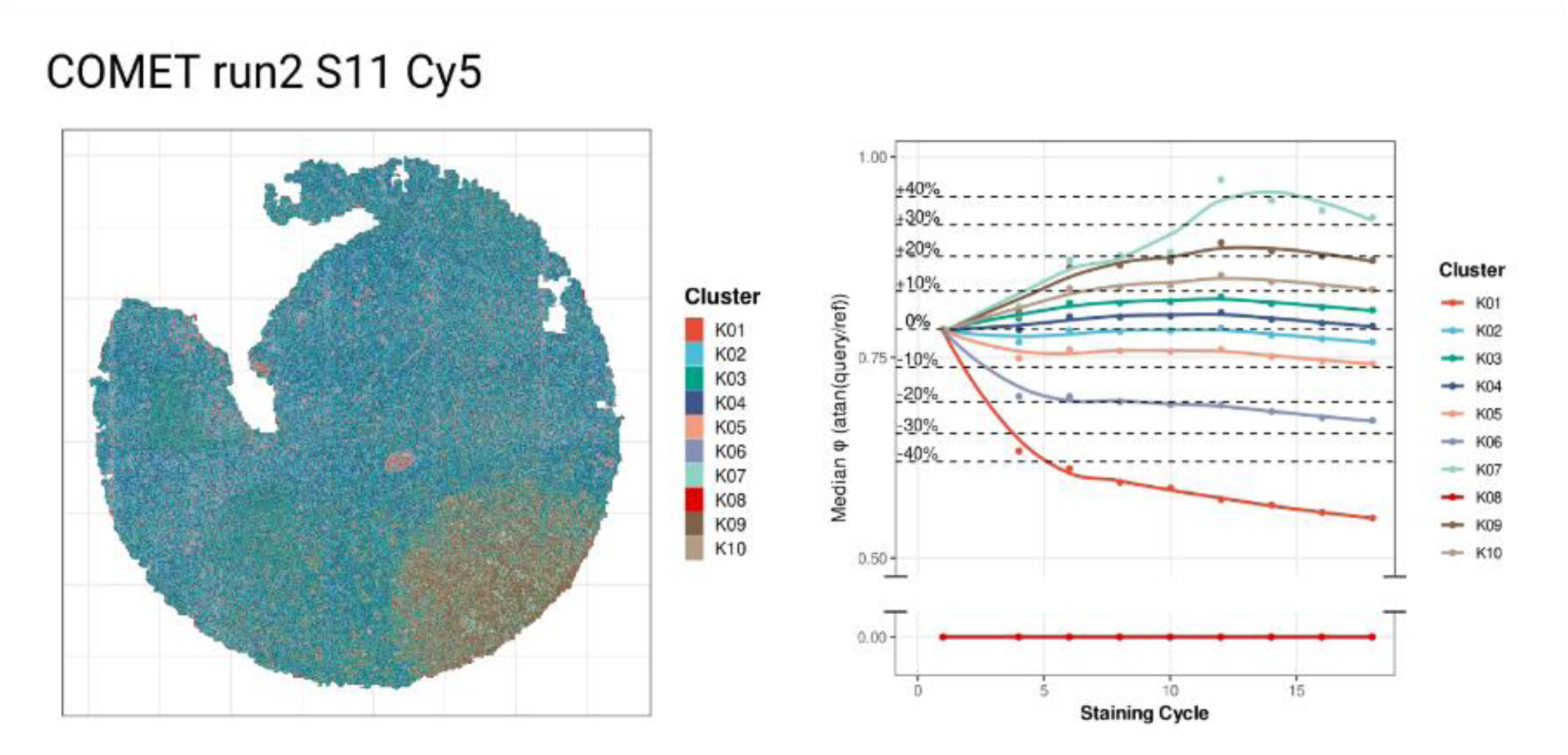
Intra-tissue signal drift across cycles. Left) Spatial cluster assignment (k=10) from baseline (cycle 1) AF image for COMET S11 (Cy5 channel). **Right)** Cluster-specific trajectories of median φ across 18 cycles. Tumour-enriched clusters (K09-K10) display increasing AF signal over cycles, whereas stromal and vascular clusters (K01-K08) show progressive intensity loss, revealing pronounced regional heterogeneity in AF dynamics.

**Supplementary Figure S4.**
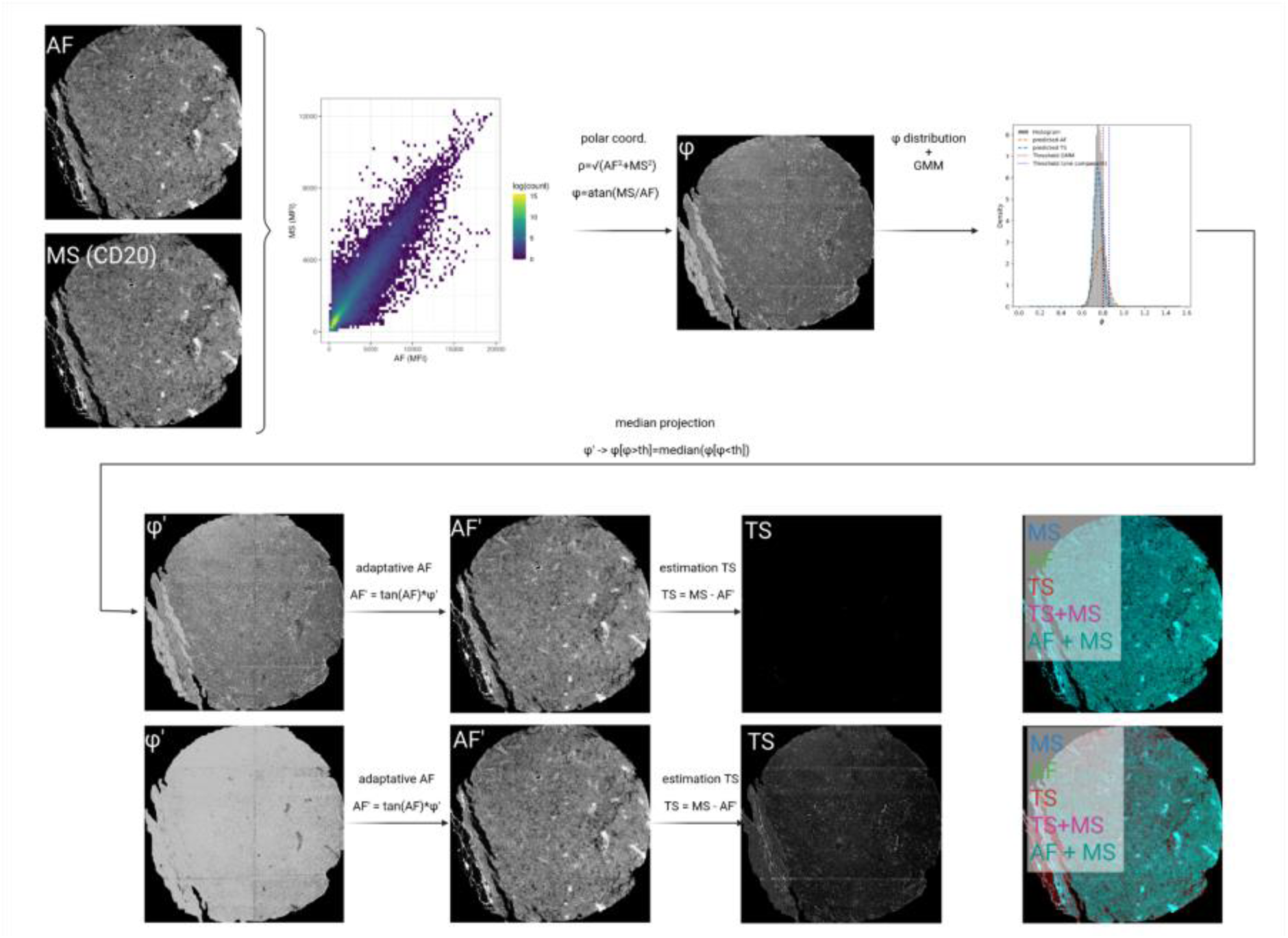
Handling low-TS images by unimodal modelling in autofluorescence subtraction. Application of CLEAR-AF on a case with minimal true signal (CD20). Left: Input images include an autofluorescence-only (AF) image and a mixed signal image (MS) acquired from the same field (CD20 channel shown). Middle: A joint intensity scatterplot (AF vs MS) is transformed to polar coordinates. In this case the φ distribution is modelled using a single Gaussian due to the absence of a clearly separable TS population. Lower branch (top): Pixels above the threshold are projected to the autofluorescence manifold using a median projection. Adaptive AF estimates (AF’) are then computed per pixel, and TS is derived via TS = MS – AF’. Right: Composite image visualizing the separate components (MS=blue, AF=green, TS=red), highlighting correct subtraction of AF (black CD20 signal). Lower branch (top): Equivalent following a two component GMM decomposition. Note how the TS image captures fluorescence not corresponding to CD20 staining (false positives).

**Supplementary Table 1. Overview of experimental design of the experiment that generated our dataset.** From left to right the table contains the cycle number, the channel in which the data was acquired, the antibody that was acquired in that particular cycle-channel and the exposure time (necessary for the NMF method). To be found in SupplementaryTable1.xlsx file.

To be found in SupplementaryTable1.xlsx file.

## References

1. Antoranz, A., Arnould, A., Andhari, M. D., Nazari, P., Shankar, G., Moor, B. De Smet, F. De Bosisio, F. M., & Pey, J. (2024). COLLAGE: COnsensus aLignment of muLtiplexing imAGEs. BioRxiv, 2024.07.15.603557. 10.1101/2024.07.15.603557

2. Antoranz, A., Van Herck, Y., Bolognesi, M. M., Lynch, S. M., Rahman, A., Gallagher, W. M., Boecxstaens, V., Marine, J. C., Cattoretti, G., van den Oord, J. J., De Smet, F., Bechter, O., & Bosisio, F. M. (2022). Mapping the Immune Landscape in Metastatic Melanoma Reveals Localized Cell-Cell Interactions That Predict Immunotherapy Response. Cancer Research, 82(18), 3275–3290. 10.1158/0008-5472.CAN-22-0363

3. Baharlou, H., Canete, N. P., Bertram, K. M., Sandgren, K. J., Cunningham, A. L., Harman, A. N., & Patrick, E. (2021). AFid: a tool for automated identification and exclusion of autofluorescent objects from microscopy images. Bioinformatics, 37(4), 559–567. 10.1093/BIOINFORMATICS/BTAA780

4. Becker, W. (2012). Fluorescence lifetime imaging--techniques and applications. Journal of Microscopy, 247(2), 119–136. 10.1111/J.1365-2818.2012.03618.X

5. Bolognesi, M. M., Manzoni, M., Scalia, C. R., Zannella, S., Bosisio, F. M., Faretta, M., & Cattoretti, G. (2017). Multiplex Staining by Sequential Immunostaining and Antibody Removal on Routine Tissue Sections. The Journal of Histochemistry and Cytochemistry : Official Journal of the Histochemistry Society, 65(8), 431–444. 10.1369/0022155417719419

6. Chevrier, S., Crowell, H. L., Zanotelli, V. R. T., Engler, S., Robinson, M. D., & Bodenmiller, B. (2018). Compensation of Signal Spillover in Suspension and Imaging Mass Cytometry. Cell Systems, 6(5), 612–620.e5. 10.1016/J.CELS.2018.02.010

7. Croce, A. C., Ferrigno, A., Bottiroli, G., & Vairetti, M. (2018). Autofluorescence-based optical biopsy: An effective diagnostic tool in hepatology. Liver International, 38(7), 1160–1174. 10.1111/LIV.13753;PAGEGROUP:STRING:PUBLICATION

8. Demchenko, A. P. (2020). Photobleaching of organic fluorophores: quantitative characterization, mechanisms, protection. Methods and Applications in Fluorescence, 8(2). 10.1088/2050-6120/AB7365

9. Eng, J., Bucher, E., Hu, Z., Zheng, T., Gibbs, S. L., Chin, K., & Gray, J. W. (2022). A framework for multiplex imaging optimization and reproducible analysis. Communications Biology 2022 5:1, 5(1), 438-. 10.1038/s42003-022-03368-y

10. Eng, J. R., Bucher, E., Hu, Z., Walker, C. R., Risom, T., Angelo, M., Gonzalez-Ericsson, P., Sanders, M. E., Chakravarthy, A. B., Pietenpol, J. A., Gibbs, S. L., Sears, R. C., & Chin, K. (2025). Highly multiplexed imaging reveals prognostic immune and stromal spatial biomarkers in breast cancer. JCI Insight, 10(3), e176749. 10.1172/JCI.INSIGHT.176749

11. Giorgino, T. (2009). Computing and Visualizing Dynamic Time Warping Alignments in R: The dtw Package. Journal of Statistical Software, 31(7), 1–24. 10.18637/JSS.V031.I07

12. Hwang, W., Raymond, T., McPartland, T., Jeong, S., & Evans, C. L. (2024). Fluorescence Lifetime Multiplexing (FLEX) for simultaneous high dimensional spatial biology in 3D. Communications Biology 2024 7:1, 7(1), 1012-. 10.1038/s42003-024-06702-8

13. Jun, Y. W., Kim, H. R., Reo, Y. J., Dai, M., & Ahn, K. H. (2017). Addressing the autofluorescence issue in deep tissue imaging by two-photon microscopy: the significance of far-red emitting dyes. Chemical Science, 8(11), 7696–7704. 10.1039/C7SC03362A

14. Karimi, E. L. H. A. M., S. N., M. N., T. W., A. A., Q. N., A. L., A. R., A. A., G. N. M. and C. N. (2024). Method of the Year 2024: spatial proteomics. Nature Methods 2024 21:12, 21(12), 2195–2196. 10.1038/s41592-024-02565-3

15. Lakowicz, J. R. (2006). Principles of fluorescence spectroscopy. Principles of Fluorescence Spectroscopy, 1–954. 10.1007/978-0-387-46312-4/COVER

16. Lin, J. R., Fallahi-Sichani, M., & Sorger, P. K. (2015). Highly multiplexed imaging of single cells using a high-throughput cyclic immunofluorescence method. Nature Communications 2015 6:1, 6(1), 8390-. 10.1038/ncomms9390

17. Monici, M. (2005). Cell and tissue autofluorescence research and diagnostic applications. Biotechnology Annual Review, 11(SUPPL.), 227–256. 10.1016/S1387-2656(05)11007-2

18. Neumann, M., & Gabel, D. (2002). Simple method for reduction of autofluorescence in fluorescence microscopy. Journal of Histochemistry and Cytochemistry, 50(3), 437–439. 10.1177/002215540205000315;PAGE:STRING:ARTICLE/CHAPTER

19. Peng, T., Thorn, K., Schroeder, T., Wang, L., Theis, F. J., Marr, C., & Navab, N. (2017). A BaSiC tool for background and shading correction of optical microscopy images. Nature Communications 2017 8:1, 8(1), 14836-. 10.1038/ncomms14836

20. Potier, G., Doméné, A., & Paul-Gilloteaux, P. (2022). A flexible open-source processing workflow for multiplexed fluorescence imaging based on cycles. F1000Research, 11, 1121. 10.12688/F1000RESEARCH.124990.1/DOI

21. Richardson, D. S., & Lichtman, J. W. (2015). Clarifying Tissue Clearing. Cell, 162(2), 246–257. 10.1016/j.cell.2015.06.067

22. Rodrigues, N. T. L., Bland, T., Borrego-Pint, J., Ng, K., Hirani, N., Gu, Y., Foo, S., & Goehring, N. W. (2022). SAIBR: A simple, platform-independent method for spectral autofluorescence correction. Development (Cambridge), 149(14). 10.1242/DEV.200545/276004

23. Sapio, M. R., King, D. M., Maric, D., Shah, S. R., Talbot, T. L., Manalo, A. P., Nara, P., Ma, W., Ghetti, A., Ramsden, C. E., Iadarola, M. J., & Mannes, A. J. (2025). Efficient removal of naturally-occurring lipofuscin autofluorescence in human nervous tissue using high-intensity white light. The Journal of Pain, 30, 105359. 10.1016/J.JPAIN.2025.105359

24. Schapiro, D., Sokolov, A., Yapp, C., Chen, Y. A., Muhlich, J. L., Hess, J., Creason, A. L., Nirmal, A. J., Baker, G. J., Nariya, M. K., Lin, J. R., Maliga, Z., Jacobson, C. A., Hodgman, M. W., Ruokonen, J., Farhi, S. L., Abbondanza, D., McKinley, E. T., Persson, D., … Sorger, P. K. (2021). MCMICRO: a scalable, modular image-processing pipeline for multiplexed tissue imaging. Nature Methods 2021 19:3, 19(3), 311–315. 10.1038/s41592-021-01308-y

25. Sun, Y., Ip, P., & Chakrabartty, A. (2017). Simple Elimination of Background Fluorescence in Formalin-Fixed Human Brain Tissue for Immunofluorescence Microscopy. Journal of Visualized Experiments (JoVE), 2017(127), e56188. 10.3791/56188

26. Tan, W. C. C., Nerurkar, S. N., Cai, H. Y., Ng, H. H. M., Wu, D., Wee, Y. T. F., Lim, J. C. T., Yeong, J., & Lim, T. K. H. (2020). Overview of multiplex immunohistochemistry/immunofluorescence techniques in the era of cancer immunotherapy. Cancer Communications, 40(4), 135–153. 10.1002/CAC2.12023;JOURNAL:JOURNAL:25233548;WGROUP:STRIN G:PUBLICATION

27. Wang, B., Li, M., Huang, X., Liu, B., Wang, B., Li, M., Huang, X., & Liu, B. (2025). Emerging trends of fluorescence lifetime imaging microscopy (FLIM): advances, challenges, and prospects. Biophysics Reports, 0(0), 0. 10.52601/BPR.2025.250024

28. Wang, S., Ren, X., Wang, J., Peng, Q., Niu, X., Song, C., Li, C., Jiang, C., Zang, W., Zille, M., Fan, X., Chen, X., & Wang, J. (2023). Blocking autofluorescence in brain tissues affected by ischemic stroke, hemorrhagic stroke, or traumatic brain injury. Frontiers in Immunology, 14, 1168292. 10.3389/FIMMU.2023.1168292/BIBTEX

29. Woolfe, F., Gerdes, M., Bello, M., Tao, X., & Can, A. (2011). Autofluorescence removal by non-negative matrix factorization. IEEE Transactions on Image Processing, 20(4), 1085–1093. 10.1109/TIP.2010.2079810

30. Zhang, D., Rubio Rodríguez-Kirby, L. A., Lin, Y., Wang, W., Song, M., Wang, L., Wang, L., Kanatani, S., Jimenez-Beristain, T., Dang, Y., Zhong, M., Kukanja, P., Bao, S., Wang, S., Chen, X. L., Gao, F., Wang, D., Xu, H., Ma, C., … Fan, R. (2025). Spatial dynamics of brain development and neuroinflammation. Nature 2025 647:8088, 647(8088), 213–227. 10.1038/s41586-025-09663-y

31. Zimmermann, T., Rietdorf, J., & Pepperkok, R. (2003). Spectral imaging and its applications in live cell microscopy. FEBS Letters, 546(1), 87–92. 10.1016/S0014-5793(03)00521-0

